# Brain Feature Maps Reveal Progressive Animal-Feature Representations in the Ventral Stream

**DOI:** 10.1101/2024.11.24.625066

**Authors:** Z. Zhang, T.S. Hartmann, R.T. Born, M.S. Livingstone, C.R. Ponce

## Abstract

What are the fundamental units of representation in the primate visual brain? While *objects* have become an intuitive framework for studying neurons in many parts of cortex, it is possible that neurons follow a more expressive organizational principle, such as encoding generic features present across textures, places, and objects. In this study, we used multi-electrode arrays to record from neurons in early (V1/V2), middle (V4), and late (posterior inferotemporal cortex (PIT)) areas across the visual hierarchy, estimating the local operation of each neuron over the entire visual scene. These estimates, called “heatmaps,” approximate the *weight sharing* operation of convolutional neural networks. We found that while populations of neurons across V1, V4, and PIT responded over the full scene, they focused on salient sub-regions within object outlines. The best captured object feature class belonged to *animals*, not general objects, as a trend that increased over the visual hierarchy. These results show that the monkey ventral stream is partially organized to encode local animal features over objects, even as early as primary visual cortex.

## Introduction

Given a natural image, we are able to perceive many of its features: from the layout of the scene to the identity of objects, their position, and the textured surfaces against which they stand. How do neurons in the brain respond to these many different features? In early studies, inferotemporal cortex (IT) neurons were described as acting in a distributed fashion, responding to general “aspects of shape, texture, or colour”^1^ lacking a simple world-based interpretation. However, it was also common to find neurons responding actively to “faces or hands”^1^, which inspired many studies of visual cortex neurons using semantic categories^2–6^. How do we reconcile object-level perception with the many neurons that participate in distributed coding, being activated by local features present across many object categories, textures, and natural scenes^7–9^? One way is to examine population activity, and one hypothesis is that although many individual neurons and local neuronal clusters (*multiunits*) may be tuned to local features, neuronal populations might be activated by the entire object. For example, in the case of a face, some neurons might become activated by the eyes, some by the nose, and others by the mouth, and their collective activity would represent the *face* configuration. Showing this is difficult, due to the inherent challenge of recording from neurons whose receptive fields span evenly across the retinal image, particularly when across multiple cortical areas. Here, we examined how neuronal populations responded across entire naturalistic scenes using a local-to-global approach. Specifically, we tested how neurons at the border of V1/V2, V4, and IT responded across entire natural scenes, by measuring their activity in response to different regions of the same natural scene. This created so-called *heatmaps*^10^, inspired by an operation in convolutional neural networks named “weight sharing,” where a given filter (*hidden unit*) is convolved across an input scene (*shared* across locations). By averaging across single- and multiunit-level neuronal heatmaps within a given area, we were able to identify visual attributes evoking the highest responses of populations in these areas and ask how neurons captured entire objects vs. backgrounds. For comparison, we also investigated this question in state-of-the-art neural networks (CNNs, CRNNs, ViTs), and related the heatmaps to algorithms that predict viewing behavior. We found that the populations of visual neurons were largely activated by local regions contained within objects, yet there was an increasing focus on animal features, such as faces and hands — a trend that best explained tuning along the primate visual hierarchy, but not across all artificial networks.

## Results

We recorded neuronal population responses in V1/V2, V4, and IT. In each session, we mapped each site’s receptive field (RF). To create a heatmap, we simulated the “weight sharing” operation using RFs from either V1/V2, V4, or IT (one population heatmap per area), as was done in a previous study^10^. Specifically, instead of moving the filter around the image as done in convolutional networks, we moved the image relative to the fixed RF: the animals performed a fixation task while a 16° x 16° natural scene image (embedded on 30° x 30° Brownian noise) was flashed at different positions within a 9° x 9° grid (spacing of 2°) relative to the center of the population RF (Fig. 1, a-c). Each scene included a variety of natural and artificial settings as well as computer-generated textures. The scenes were shown for 100 ms ON and 100 to 200 ms OFF, with 3-14 flashes between juice rewards; different scenes were interleaved within a trial (**Fig. 1c**). We measured the neuronal firing rate for 200 ms, from 50 ms before image onset to 50 ms after image offset. The raw measured output at each time step was a set of firing rates arranged into a 2-D grid (a heatmap), where each row and column indices corresponded to the location of the scene, and the value was the firing rate evoked by that part of the scene (**Fig. 1d, e**). These heatmaps were resized to the original stimulus scenes (16° x 16°) and superimposed for visual inspection (**Fig. 1f-g**). The resulting heatmap revealed the neurons’ preferred shapes in the scenes, highlighting the specific parts they were most activated by, eliminating investigator bias and stimulus pre-processing. We used six monkeys with chronic microelectrode arrays in V1/V2, V4, and IT (two monkeys in all three areas, one monkey in posterior IT only, and three monkeys in V1 only), with up to 64 channels total per monkey (**Fig. 1h**).

**Fig. 1.**
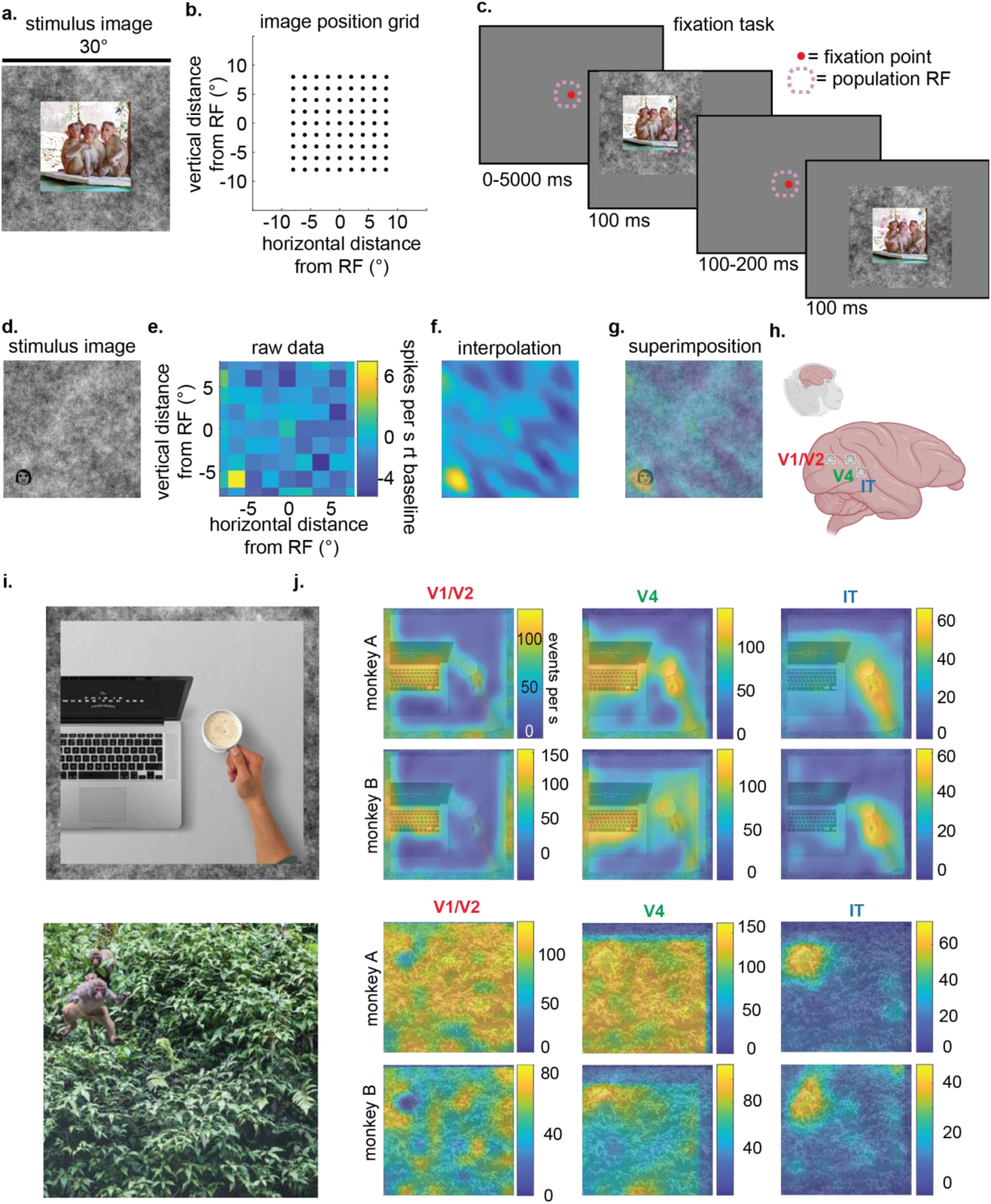
Heatmaps. **a.** Example of stimulus picture: a 16° x 16° natural scene embedded in a 30° x 30° brown noise background. **b.** Grid of 81 positions for each stimulus presentation, relative to the center of the population RF at (0,0)°. **c.** The task: monkeys fixated a central spot (red) while stimuli were shown for 100 ms over the RF (dashed pink outline). **d-g.** Simplest case of an observed feature map. **d.** The stimulus contained a 2°-diameter cartoon face in a noise background. **e.** Spike rate response elicited by every part of the picture over the 81 positions: this is the feature map. **f.** Feature map resized and interpolated (via `imresize.m`, MATLAB) to 16°x16° and **g.** superimposed on the stimulus. **h.** Schematic showing approximate location of implanted chronic arrays. **i.** Examples of natural scenes used as stimuli. **h.** Averaged heatmaps of neuronal populations in V1/V2 (column 1), V4 (column 2) and IT (column 3) in monkey A and monkey B generated from the natural scenes shown in **i.** The heatmaps may show a gap between the edge of the image: this reflects correction for the center position of the population RF, as these differed as a function of visual area.

### Neurons across the hierarchy showed increasing focus on animals

Every single- and multiunit, *j_a_*, in an area array *a* ϵ {*V*1/*V*2, *V*4, *IT*} generated one heatmap *H_j_a__*(*i*) for a natural scene *i* ϵ *I* (**Fig. 1i**), corresponding to the time-averaged activity over the first 200 ms after scene onset. To assess areal differences in tuning, we normalized each site’s heatmap *H_j_a__*(*i*) to the range 0–1 and examined the heatmap *H_a_*(*i*) averaged over *c* channels per array.

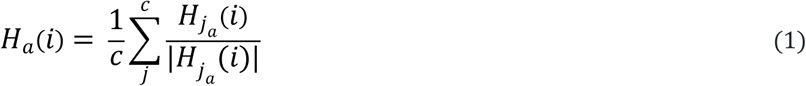

We found that within each area, the neuronal population responded to every part of the scene, but some regions in the scene evoked stronger activity than others. In particular, we found that V1/V2 neurons mainly responded strongly to backgrounds and spatial-frequency regions, IT neurons primarily responded strongly to animal features, and V4 neurons responded to both. For example, when presented with a scene containing a laptop computer next to a hand holding a cup, V1/V2 neurons responded more to the black and white keyboard — a pattern with repetitive arrangement of keys, and less to the hand holding the cup — a human body part. In contrast, IT neurons responded more to the hand holding the cup, and less to the keyboard. V4 neurons showed responses to all of these objects (**Fig. 1j, rows 1-2**). As a second example, when presented with a scene of a distant macaque jumping in front of a rich green canopy, V1/V2 neurons responded more to the canopy, IT neurons responded more to the monkey, and V4 neurons responded to both (with a subtle emphasis on the macaque, **Fig. 1j, rows 3-4**). Generally, we found that deeper along the ventral pathway, population responses signaled animal features, such as monkey and human faces, hands, and bodies in the natural images. However, it is important to reiterate that neuronal populations *responded across the whole scene*, but populations differed in the specific regions that they emphasized overall.

While this could signal a shift in tuning towards objects generally, the fact that a *keyboard* evoked less population activity than a *hand* (both being “objects”) suggested that more elemental features were behind this response. To quantify how the responses of different areas changed based on the presence of object type and shape, we obtained an independent assessment of the scene’s visual content. Specifically, we created binary masks *M*(*i*) representing different information within each scene *i*, which we could use to measure how the neuronal heatmaps were contained within each mask. We computed these maps independently of neuronal data, using both algorithmic and crowd-sourcing approaches, and obtained a total of 27 possible semantic masks per scene (**Fig. 2a, Methods**). These masks linked specific scene regions with object labels. After obtaining a set of masks per scene, we merged masks of subordinate labels into single masks of superordinate categories (such as “moss” into “plants,” or “snouts,” “monkey” etc. into “animals”). Next, we superimposed each mask *M*(*i*) over the heatmaps, and measured the mean neuronal response over time inside the mask *R_in,a_*(*i, t*) and outside the mask *R_out,a_*(*i, t*), for V1/V2, V4, and IT (**Fig. 2b**) as

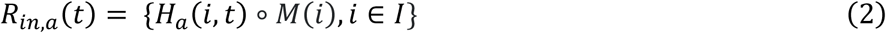

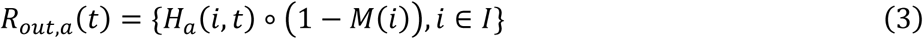

where *t* was time, as each heatmap was first computed every 10 ms over the 200 ms response time. We first examined how the *Animal* mask captured the responses of these areas. We found that the mean activity difference between *R_in,a_*(*t*) and *R_out,a_* (*t*) was the largest in IT, followed by V4, with the smallest difference in V1 (**Fig. 2c**, top). We quantified this effect by plotting the difference *D_a_*(*t*) = *R_in,a_*(*t*) − *R_out,a_* (*t*) and measuring the area under the curve (AUC) of *D_a_*(*t*). As a key control, we shuffled the response regions for each scene to obtain a shuffled heatmap *H_a_(shufled)__*(*i, t*) and also measured the AUC for *D_a_shuffled__* (*t*).

**Fig. 2.**
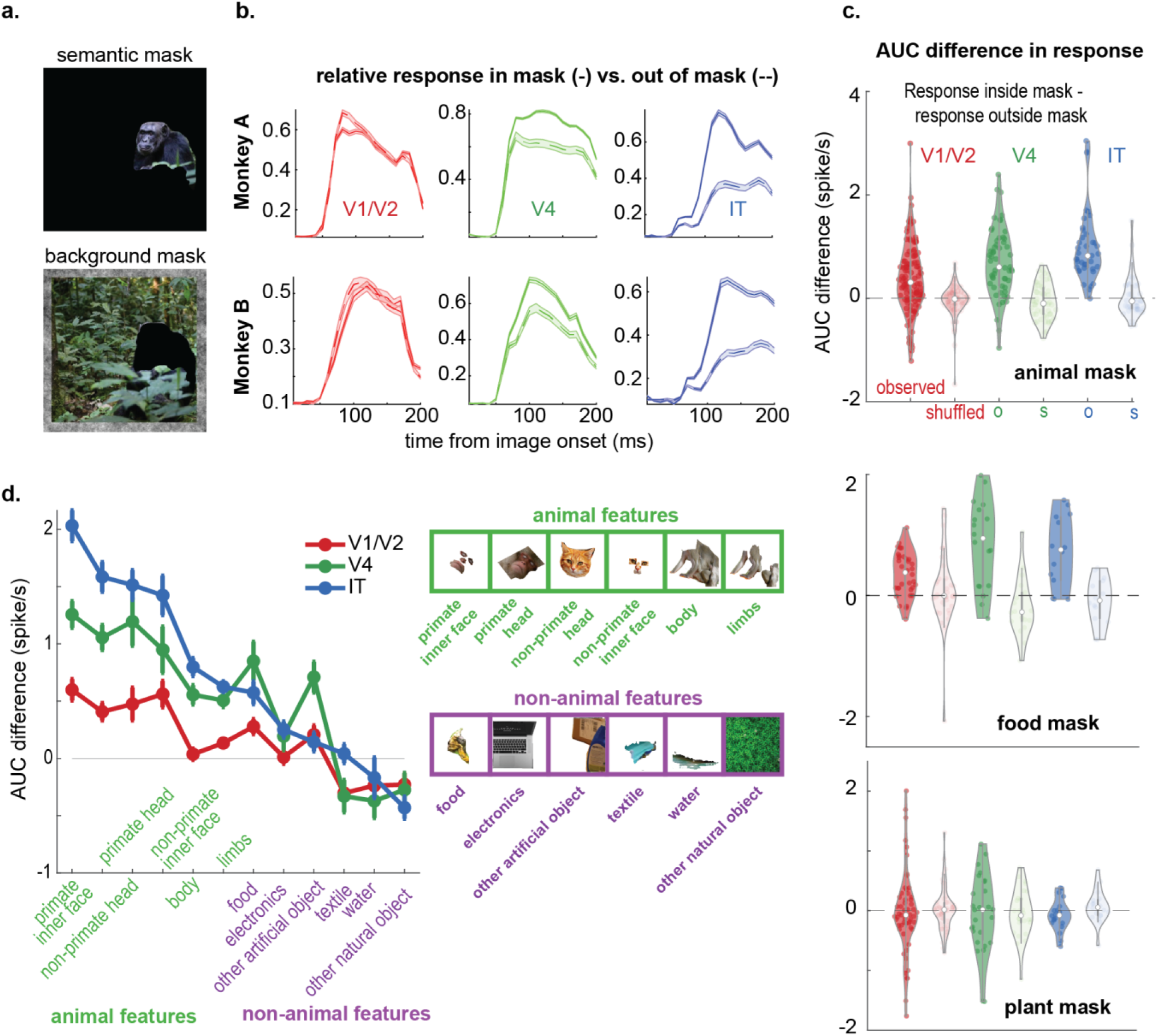
Neurons across the hierarchy showed increasing focus on animal features. **a.** Example of the *Animal* masks derived from the scene of a chimpanzee in a jungle (top) and the background mask of the same image. **b.** Mean (population heatmap) activity inside (solid lines) and outside of the mask (dashed lines, ±SEM), for V1/V2 (red), V4 (green) and IT (blue), plotted as a function of image onset. **c.** Aggregate difference between the curves, measured as the area under the curve (AUC) for the difference curve defined by the in- vs. outside response curves *AUC*(*D_a_*(*t*)), for V1/V2, V4 and IT (red, green, blue, dark colors, *N =283*, comprising responses of neuronal populations for six animals, up to 36 pictures); the observed values in dark colors are compared to the shuffled-map AUC values (lighter colors). Top plot: *Animal* mask, middle plot: *Food* mask, bottom plot: *Plant* mask. **d.** Time averaged neuronal response E[*AUC*(*D _a_*(*t*))] per visual area, across all animals, for representative categories. Insets show examples of the content of the animal masks.

*Animal* masks identified regions evoking stronger neuronal activity for IT than for V4, and stronger for V4 than for V1. Still, these masks isolated V1 responses better than they did for shuffled controls (**Fig. 2c, top**, observed AUC median ± SE; V1/V2: 0.19 ± 0.01, V4: 0.63 ± 0.06, IT: 0.73 ± 0.08; shuffled maps: V1: −0.0083 ± −0.0002, V4: 0.005 ± 0.004, IT: −0.030 ± −0.001). This suggests that the *Animal* masks represent a mix of low- and high-level saliency features, such as those detected by shallow algorithms like graph-based visual saliency metrics^11^, but also by deeper models such as CNNs. To capture this change across areas, we used a linear model to measure the slope across V1/V2, V4, and IT AUC values. Higher slope estimates indicate an increase in *within-mask* neuronal response from V1/V2 to IT. Then, we asked if we could identify the trend in neuronal responses to other categories such as *Food* and *Plants*, in addition to (subordinate) categories.

Generally, the slope values for *Animal* masks were higher than for non-*Animal* masks (**Fig. 2c, middle and bottom**). Specifically, the highest slope values were for primate *Inner*- and *Outer*-*Face*, followed by non-*Primate Inner*- and *Outer*-*Face*, and then by *Body*, *Limbs*. Slopes decreased further by non-*Animal* masks, across *Food*, *Textiles*, *Electronics*, and *Water* (as examples, separately considering *food, books, environment, plant,* and other labels; AUC values, median ± SE; V1: 0.022 ± 0.003, V4: 0.16 ± 0.01, IT: 0.006 ± 0.003; AUC for shuffled maps: V1, 0.0057 ± 0.0003, V4: −0.031 ± 0.002, IT: 0.002 ± 0.002; **Fig. 2d**). These results suggest that the monkey visual ventral pathway, as early as V1, is organized, at least in part, to increasingly encode background-isolated regions containing animal features. The fact that results differed as a function of mask content suggests this trend was not a trivial consequence of the analysis or from architectural interareal differences such as receptive field size. One important caveat is that the parts of the lateral ventral stream from which we recorded emphasize the central visual field; a different set of representational patterns might be found when recording from neurons representing eccentric regions of the visual field, as these neurons show tuning for less curved features^12,13^.

### Populations responded to objects, but prioritized local features

Above, we showed that areas across the ventral stream progressively prioritized the processing of animal information, with the strongest focus observed in IT. Conversely, areas like V1 and V4 showed strong processing activity to more background and inanimate visual features. Upon visual inspection, we observed that the neuronal heatmaps highlighted certain subregions within the broader semantic masks, instead of fully encompassing the entire object outline as defined by human participants and object-trained deep networks. To delve deeper into this observation, we further investigated whether neuronal populations processed *category* information by selectively responding to localized visual features, to the overall segmented object, or to a combination of both.

For this analysis, we used two monkeys with arrays implanted in V1/V2, V4, and IT (monkeys A, and B). To quantify the spatial extent of neuronal activity, we introduced a metric called a *feature-to-object ratio* (FTO), an estimate of the neuronal activity’s coverage within an object. Here, *objects* are defined as semantic classes that could be outlined by masks *M_o_*(*i*) of object *o* of scene *i*, such as *monkey* or *car* (**Fig. 3a**). If the peak neuronal activity were uniformly distributed within the object, the FTO would attain a value of 1 (**Fig. 3b**); conversely, if none of the peak responses fell inside the object outline, the FTO would be 0. To compute the FTO, we first normalized the neuronal activity heatmaps *H_a_*(*i*) to the range of 0 and 1, so that the FTO of a scene *i* in array *a* was obtained by summing the normalized responses within the object mask and dividing by the sum of the identity matrix within the object mask. To control the relative frequency of response hotspots within a scene, we shuffled the activity within the map, as in the above analyses.

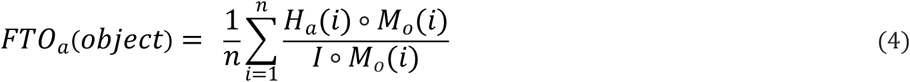

**Fig 3.**
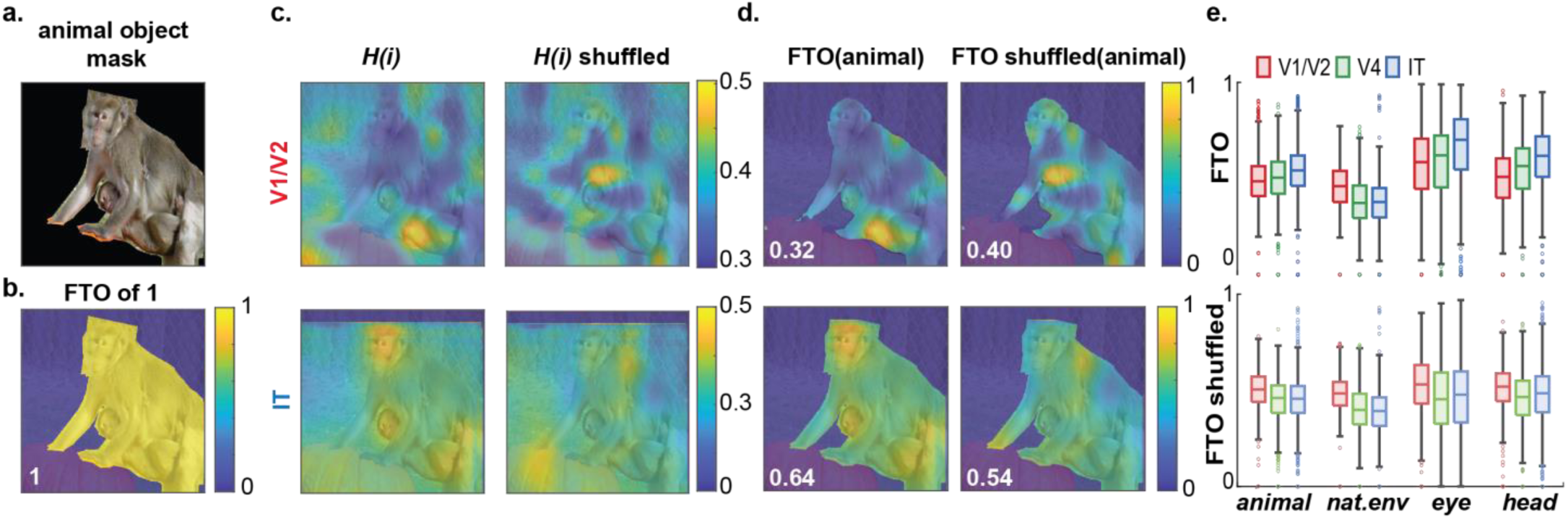
Feature to Mask. **a.** animal mask of a natural scene of a mother monkey and a baby monkey. **b.** Illustration of neuronal activity was distributed evenly within the object outline resulting in FTO of 1 **c.** IT and V1 population heatmap and shuffled heatmap in Monkey A. **d.** FTO(animal) of IT and V1 heatmaps in b. **e.** Top: FTO ratio of Monkey A and B in V1/V2 (red), V4 (green), IT (blue) for animal object); the observed FTO values in dark colors are compared to the shuffled-map FTO values (lighter colors). Bottom: FTO for natural texture object.

We found that the FTO ratio was less than 1 for all visual areas. We also found that the median FTO ratio to *Animal* masks depended on visual area, increasing from V1/V2 to IT (V1/V2, V4, IT values median ± SE, Monkey A: 0.5144 ± 0.0001, 0.5295 ± 0.0002, 0.5514 ± 0.0001, ANOVA: F=5.713, P=3.4×10^-3^, DF=2,2169); Monkey B: 0.4493 ± 0.0002, 0.4845 ± 0.0003, 0.5286 ± 0.0002; ANOVA: F=25.305 p=1.5×10^-11^, DF=2,1747) (**Fig. 3c,d**). We fit a linear model to the median FTO ratio across V1, V4, and IT and compared the slope *k*. The trend across visual areas depended on specific categories and segmentation mask size: for example, the median *k*(FTO(*head*)) was larger than for *k*(FTO(*animal*), which was larger than *k*(FTO(*natural texture*)) (**Fig. 3e**). This result suggests that neuronal populations ranging from V1/V2, V4, and posterior IT were more turned to specific localized features within objects than to the overall identity of the animal objects.

### Some deep networks showed an animal-feature focus

Next, we asked if any state-of-the-art visual hierarchical system developed a focus on animal features as the primate visual system did. We searched for this phenomenon in artificial neural networks (ANNs), selecting 18 models with architectural diversity, including convolutional neural networks (CNNs), convolutional recurrent neural networks (CRNNs), and vision transformers (ViTs) pretrained on ImageNet^14^. We also wanted to assess whether a model’s capability to match neuronal response patterns could anticipate this phenomenon. To achieve this, we ensured that the selected models displayed a diverse range of scores in the Brain-Score benchmark^16^. To simulate the neurophysiological experiments above, we tested each model with the same set of scenes *I*, and sampled three layers at different depths in each artificial model (“early,” “middle,” and “late”) to emulate the three cortical stages (V1/V2, V4, PIT). Once each layer was selected, we randomly sampled *c* = 32 channels within that layer, simulating the sampling of the microelectrode arrays (**Fig. 4a**, see **Methods**). For each ANN layer *l*, we defined the averaged normalized activated feature map to natural scene *i* as *F_l_*(*i*):

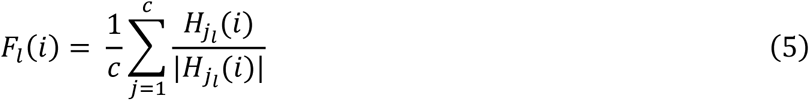

**Figure 4.**
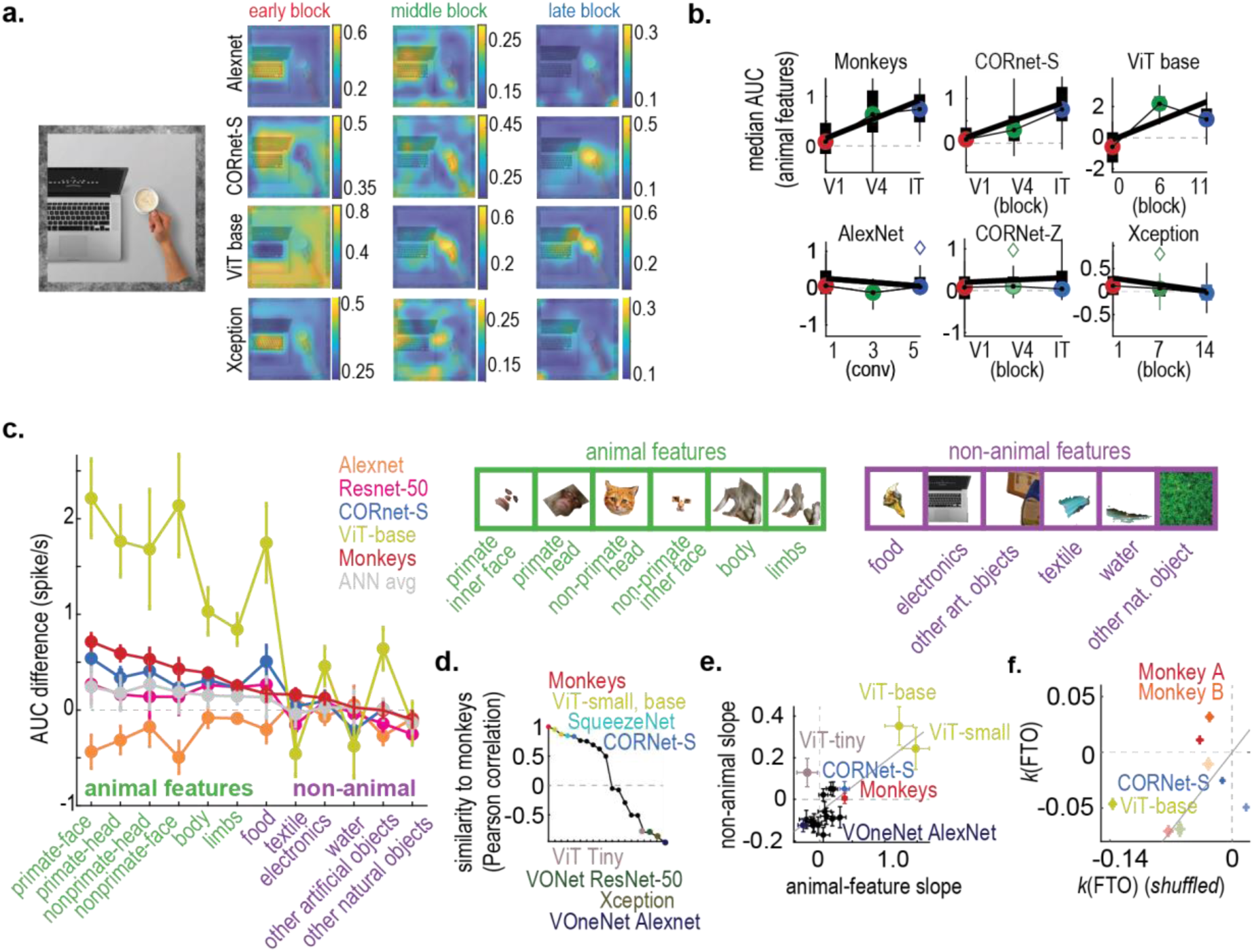
Evaluation of Deep Neural Networks on Brain Similarity. **a.** Example of the heatmaps from monkeys from V1 to V4 to IT and ANNs in early, middle, late layers to the scene of a person with partial face visible **b.** Mean (population heatmap) activity inside (solid lines) and outside of the mask (dashed lines, ±SEM), for V1/V2/early block (red), V4/middle block (green) and IT/late block (blue) **c.** non-animal features slope vs animal features slope for monkeys and ANNs **d.** Pearson correlation measuring of ANNs similarity to Monkeys **e.** slope of non-animal feature vs animal features for monkey and ANNs. **f.** slope of FTO vs slope of FTO in control for animal features (dark colors), and natural texture feature (lighter colors) in Monkeys, CORnet S and ViT base.

As in the *in-vivo* experiments, we quantified mean activation inside the mask *R_in,l_* and the mean activation outside the mask *R_out,l_* for each layer *l* in every model *m*. We then measured the area under the curve *D_l_* = *R_in,l_* − *R_out,l_* (AUC) (**Fig. 4b, Supplementary Table A**).

We found that as a group, ANNs showed a wide diversity of behaviors. Heatmaps in some ANNs such as robust ResNet-50 and CORNet-S displayed the same increasing focus toward animal features, whereas other ANNs such as Xception and AlexNet did not. To quantify if an ANN trend was similar to the monkey trend, we used three criteria: (1) the ANN should show an increasing focus on animal features across layers, (2) this increasing trend would be of comparable magnitude to the neurophysiology data, and (3) not only with *Animal* features but also subordinate categories (**Fig. 4c, Supplementary Table B**). We used a series of tests to narrow the list of ANNs that act more like the monkey brain.

To test criteria (1), we measured the ordinary least square (OLS) fitting slope *k_a_* to animal-features across *D_l_early__*, *D_l_middle__*, *D_l_late__* for each monkey or model *m*, and also the slope *k_na_* to nonanimal-features (**Fig. 4b, e**). Monkeys showed a positive slope across areas for *Animals* but a flat slope for *Non-Animals* (OLS slope median ± SE: *k_a_*: 0.38±0.03, *k_na_*: 0.07±0.05), indicating an increasing focus on animal features from V1/V2 to IT. CORNet-S showed a positive trend that was similar to the monkey data (*k_a_*: 0.48 ± 0.06, *k_na_*: 0.05 ± 0.04), whereas ViT-base, ViT-small, and SqueezeNet also showed positive trends but were less similar to the monkey brain (ViT-base, *k_a_*: 0.98 ± 0.14, *k_na_*: −1.33 ± 0.25; ViT-small, *k_a_*: 0.67 ± 0.07, *k_na_*: 0.15 ± 0.04; SqueezeNet, *k_a_*: 0.26 ± 0.06, *k_na_*: −0.13 ± 0.04). ANNs such as VOne-CORNet-S, Xception, and AlexNet showed flat trends to both *Animal*- and *Non-Animal* features (VOne-CORnet-S, *k_a_*: 0.08±0.08, *k_na_*: −0.11±0.05; Xception: *k_a_*: −0.15±-0.04, *k_na_*: −0.10±0.02, AlexNet *k_a_*: −0.09±0.06, *k_na_*: −0.11±0.03).

Next, we compared the trend magnitude between the ANN- and neurophysiology results by asking if there were similar effect differences *R_in, l_* between *D_l_early__*, *D_l_middle__*, and between *D_l_middle__* and *D_l_late__*. To do this, we combined two-sample tests of significance, a correlation of stage difference, and a multivariate ANOVA (MANOVA). Monkeys showed a difference *R_in, l_* between *D_l_early__*, *D_l_middle__* and between *D_l_middle__*, *D_l_late__* (two sample t-test, p value: 5.98×10^-9^, 5.72×10^-2^, DF: 195, 143). Among the ANNs that met criteria (1) with positive *k_a_* values, only VOneNet-Resnet50-at showed a significant change between both *D_l_early__*, *D_l_middle__* and *D_l_middle__*, *D_l_late__* (two-sample Student’s t-test, *P =* 1.07×10^-2^, 1.88×10^-4^, DF, 58), while respectively, Resnet-50 robust, ViT-small, CORNet-S, SqueezeNet, Resnet-50 and ViT-base showed more similar trends as the monkeys (MANOVA predictor *P* values, Resnet-50 robust: 2.2×10^-8^, ViT-small: 5.8 ×10^-12^, CORNet-S: 0.01, Squeeze Net: 2.2 ×10^-5^, Resnet-50: 3.2 ×10^-10^, ViT-base: 5.2 ×10^-6^. We also ranked each network by its similarity to the monkey trends, using a Pearson correlation coefficient, and found that the best correlations were shown by ViT small (0.96), SqueezeNet and ViT base (0.88), and CORNet-S (0.82). Among the non- and anti-correlated networks were VOneNet-ResNet-50 (−0.09), AlexNet (−0.59), and ViT tiny (−0.83, **Fig 4d**). We found a correlation between a network’s trend to capture animal features over its hierarchy, and the same network’s trend to capture non-animal features (linear fit slope = 0.24 (0.14-0.33, 95% CI), suggesting that part of the trend can be explained by a focus towards objects in general (**Fig. 4e, Supplementary Table C**). CORnet S and ViT base had a negative *k*(FTO) for both animal-and natural texture features, in contrast to positive *k*(FTO) for animal feature in monkeys (**Fig. 4f, Supplementary Table D**).

To understand why some networks matched the brain better than others, despite their large differences in architectural design, we analyzed their robustness on parametric image degradations (such as style, color, and contrast manipulations), and eventually, we narrowed our attention to model performance on texture-shape cue conflict, such as an object with airplane shape but a bear-like texture^15^ (**Supplementary Table E**). We measured the percentage of the decision of each model in classifying the image based on shape and texture. We found CORNet-S has higher classification accuracy on images with animal objects than non-animal objects (classification accuracy: 86.08%, 65.07%). In addition, CORNet-S was more biased towards textures when the images contained animals. Specifically, CORNet-S made 75.5% of classification decisions based on the object’s texture if the image contained animal objects, but this fraction of texture-based decisions was only 46.52% on images containing non-animal objects. While ImageNet-trained ANNs are known to be biased toward textures^15^, a recent study has also shown a texture bias in primate IT neurons^16^, coinciding with the strong bias towards animal texture found in CORNet-S. This suggests high selectivity to animal texture might be critical in the visual information processing in primate brain. Overall, we conclude that the majority of ANNs did not show the same pattern of monotonic increase towards animal features as the monkey brain did, and those that did partially depended on a bias towards animal-related textures.

### Free viewing and automated saliency maps correlated with V4 and IT heatmaps

These activity heatmaps showed the activity of a subpopulation of neurons over a large visual scene. One hypothesis is that the animal might be more likely to make saccades to regions of the scene that also drive the mean population activity strongly. To determine if these heatmaps could serve as a basis for bottom-up saliency, we measured how monkeys looked at the visual scenes under free-viewing conditions, and composed *fixation maps* which we could correlate with the neuronal heatmaps. We measured the similarity of neuronal heatmaps to fixation maps and to predictions made by each artificial model.

In the experiments, we measured the monkeys’ eye movements in a free-viewing task, using the same natural scenes *I*, presented at the same size as in the neuronal experiments. In this task, each monkey initiated a trial by fixating a small (0.2°-diameter) round target in the center of the monitor. After acquisition, the scene was flashed on for one second and the animal was then free to view any features in the scene. If they kept their gaze within the natural scene, they received a liquid reward. By tracking their gaze over the stimuli, we were able to create an eye-position heatmap per scene (fixation maps, **Fig. 5a-c**) and compared it with the corresponding neuronal activity heatmaps from different areas (**Fig. 5c, *ii***) for the same scenes. Finally, we processed the same visual scenes *I* using saliency models such as Graph-based visual saliency (GBVS^11^), Itti-Koch saliency^17^, and *Fast, Accurate, and Size-Aware* salient object detection (FASA implemented in DeepGaze^18^, Fig. 5c, *iii*). We correlated all fixation- and saliency maps to each other and to the V1/V2, V4, and IT activity heatmaps, both observed and shuffled (**Fig. 5d**).

**Fig. 5.**
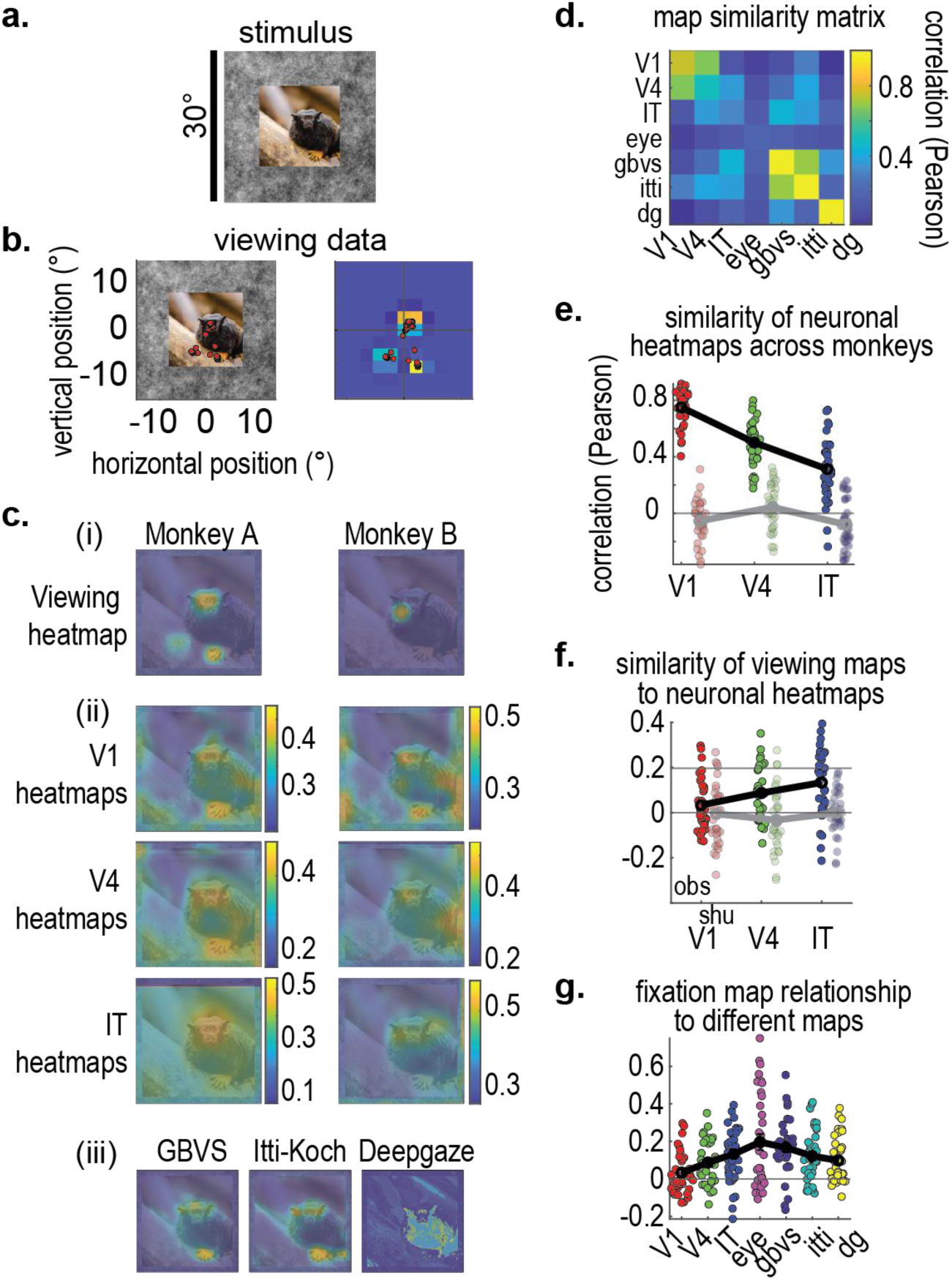
Free viewing and automated saliency maps. **a.** Example scene. **b.** Example of eye positions tracked over the scene to create fixation maps. **c.** Examples of viewing maps on the same scene (*i*), along with neuronal heatmaps from V1, V4, and IT (*ii*), and algorithmic saliency maps (graph-based visual saliency, Itti-Koch saliency, and fast, accurate, and DeepGaze-FASA, *iii*). **d.** Pearson correlation values relating all neuronal heatmaps (*V1, V4, IT*), all fixation maps (*eye*), and all saliency maps (*gbvs, itti, dg*). The diagonal cells show the monkey-monkey correlations in neuronal heatmaps and the monkey-monkey correlations in fixation maps. **e.** Similarity of neuronal heatmaps across monkeys for V1 maps, for V4 maps, and for IT maps (observed values; light red, light green, blue; shuffled neuronal heatmaps, dark red, green, and blue); each point shows one scene. **f.** Similarity of neuronal heatmaps to fixation maps, per area. **g.** Similarity between fixation maps to neuronal heatmaps and to saliency algorithms.

We found that, across monkeys, the V1 heatmaps were most closely correlated to each other (Pearson coefficient, 0.751±0.022) than were the V4 heatmaps to each other (V4: 0.670±0.026) or the IT heatmaps (IT: 0.099±0.037, **Fig. 5e**); shuffled maps showed much lower correlations (shuffled V1: −0.054±0.025, shuffled V4: 0.028±0.028, shuffled IT: −0.017±0.020). The V1 maps resembled the V4 maps more than the IT maps (Pearson correlation between V1–V4: 0.67±0.03; between V1–IT, 0.10±0.04, there was a probability of 2.5×10^-7^ these estimates arose from the same distribution (Wilcoxon signed rank test, Z-value = 5.159, N = 36). No such area-to-area proximity was evident in the shuffled maps (shuffled V1–V4, 0.03±0.03, shuffled V1–IT, −0.02±0.02, P = 0.16).

Next, we examined the fixation maps. The monkeys showed some correlation in their fixation maps, as compared across scenes (Pearson coefficient of 0.20±0.04, *P* = 5.8×10^-4^, Wilcoxon paired sign rank test, Z-value = 3.44, N = 36). Compared to the neuronal heatmaps, the fixation maps showed the weakest correlation to V1 heatmaps (0.03±0.02), an intermediate correlation to V4 maps (0.09±0.020), and a higher correlation to IT maps (0.13±0.03); there was a reliable statistical difference in this trend (*P* = 6.2×10^-3^, one-way ANOVA, area as factor, F = 5.3, d.f. = 2, N = 104,, **Fig. 5f**). There was no statistical difference between the median correlation between observed V1–fixation maps vs. the median correlation between shuffled V1– fixation maps (Pearson correlation for observed was 0.03±0.02 vs. shuffled −0.01±0.02, *P =* 0.18, Wilcoxon sign rank test, Z-value = 1.34, N = 36). However, there were statistical differences between observed V4–fixation maps vs. shuffled V4-fixation maps (0.09±0.02 vs. −0.03±0.02, *P* = 6.6×10*-4*, Wilcoxon sign rank test, Z-value = 3.41, N = 36) and between observed IT-fixation maps vs. shuffled IT-fixation maps (0.13±0.03 vs. −0.00±0.02, *P* = 3.8×10^-4^, Wilcoxon sign rank test, Z-value = 3.55, N = 36). The fixation maps also showed a reliable higher to the algorithmic saliency maps, with a stronger correlation to GBVS (0.17±0.03, *P* = 6.7×10^-6^, Wilcoxon sign rank test, Z-value = 4.50, N = 36), then to the Itti-Koch maps (0.12±0.02, *P* = 5.2×10^-5^, Wilcoxon sign rank test, Z-value = 4.05, N = 36), and lastly to the DeepGaze-II maps (0.10±0.02, *P =* 1.4×10^-4^, Wilcoxon sign rank test, Z-value = 3.82, N = 36, **Fig. 5g**). IT maps were generally better correlated to all saliency maps than were V4 and V1 maps (IT-saliency maps: 0.34±0.03, V4–saliency maps: 0.24±0.03, V1-saliency maps: 0.13±0.02, all values statistically different per one-way ANOVA with area as single factor, *P* = 3.8×10^-8^, F = 18.1, d.f. = 2, N = 314). This suggests that visual saliency algorithms overall simulate operations of more anterior visual cortex. In conclusion, we found a relationship between the monkeys’ viewing behavior and their neuronal activity heatmaps. The monkeys’ viewing patterns were more closely correlated with anterior visual cortex area heatmaps, and the same anterior visual cortex area heatmaps were also more correlated with all artificial saliency maps compared to posterior visual areas.

## Discussion

Primates perceive and interact with objects, but how is this ability reflected in neuronal populations subserving vision? While the notion that the function of individual neurons maps to object classes may seem intuitive, the sheer diversity of objects suggests that a more efficient representational scheme is necessary, one that is distributed across shared local features^1^. Recent studies suggest that most visual cortex neurons represent lower-level features: when exposed to stimuli encompassing very large image spaces, such as those represented by deep generative networks, neurons show wider response ranges^19^ and expansive tuning functions^20^, all seemingly driven by specialized local features^9^. This aligns with early investigations^7^ and is consistent with the view that objects and places must be represented in a distributed fashion across neuronal populations, if only because under most natural conditions, objects such as faces are larger than most neuronal receptive fields, even for neurons in anterior regions. There may be higher-level representations by neurons in the hippocampus, thalamus, or prefrontal cortex, but this remains to be shown using contemporary technical approaches that systematically deconstruct visual inputs^9,19^.

However, the nature of this distributed function remains unclear. Given that neurons show evoked activity in response to nearly any kind of image (while reserving their stronger activity to particular image features), one might suppose that our perception of a full scene depends on neuronal populations being systematically excited by all features in the image. Alternatively, it is plausible that perception is constructed from sparser cues, akin to how visual information in the cortical representation of the blind spot is interpolated by neighboring visual features^21^. Under this view, the role of the visual system is not to explicitly represent objects or other high-level concepts, but to simply provide a *good-enough*, “pragmatic” ensemble of features that allow interactions with the world, from navigation to grasping and recognition^22^.

In this study, we used a technique inspired by weight sharing in CNNs, where a given RF can be convolved across a full visual scene. We found that local neuronal populations (consisting of single units, multiunits, and hash) did respond to the entire visual scene, but often burst more frequently to more specific regions, depending on the visual area. For example, while V1 was driven widely by textured backgrounds, IT neurons responded to features that signal animals or objects belonging to *animate*^4^ sub-categories. While it is a well-known fact that monkey IT neurons respond strongly to face and hands^1^, many studies using these *animate* stimuli have been interpreted as evidence that these neurons respond to these objects, without localizing the features that defined these objects. Here, we found a hierarchy-wide organizing principle that informs us about the content of earlier cortical stages, such as V4 and V1/V2. Our results show a gradually increasing tendency for neural responses to emphasize the representation of features *contained* within monkey and human faces, hands, and bodies, but not covering the entire area defined by the semantic definition (spatial mask) of these objects.

This focus on animal features is consistent with the fact that macaque monkeys are foraging and social animals that spend a significant fraction of their time identifying conspecifics, classifying their actions, and avoiding threats from other animals^23,24^ (including humans in laboratory settings). In the visual world, conspecifics and predators can be occluded by trees and rocks, so monkeys often need to be able to identify other animals relying on the patches of textures perceived, instead of the whole silhouette. High curvature is a likely low-level feature prevalent in these patches, present as the outline of eyes, faces, and fingers^25,26^, but not as common in parches of artificial and other natural objects and textures. Face features are highly decodable and separable^25^, thus the focus on animal features may be directly related to curvature tuning, particularly given the lateral position of our recordings.

Detecting animal features is a problem that is also faced by deep artificial networks of vision trained for classification. ANN capability is largely dependent on their training dataset. While datasets such as ImageNet^14^ offer a diversity of scenes to simulate the natural environment, they are still far short in matching the rich streams of complex information that we experience in our daily lives. We found that only a few ImageNet^14^-trained ANNs come to share this similarity with the visual system, in learning animal-like textures, such as fur and color patterns, suggesting this to be a reliable mechanism for detection, perhaps as much as shape.

One major limitation of this study is that while weight sharing is a useful way to estimate how the visual scene is represented, a gold-standard experiment would see the collection of responses from a larger population of neurons, with receptive fields tiling the entirety of the visual scene. In principle, such a study could reveal subpopulations of neurons that respond equally strongly over the full area of a semantic mask, across variations in the semantically defined object’s size, position, and rotation. While our opinion is that this is not a necessary feature of a distributed code for object recognition, this remains to be seen. High-density recording probes, such as Neuropixels^27^, might allow for the realization of this experiment in the near future.

## Acknowledgements

We thank Mary Carter and Elizabeth Cleaveland for their technical help with data acquisition and surgical procedures.

## Author contributions

*Conceptualization:* Z.Z., T.S.H., C.R.P.*, Methodology:* Z.Z., C.R.P. *Experimentation, Visualization, Investigation:* T.S.H, Z.Z., C.R.P. *Resources and Investigation:* M.S.L, R.T.B., C.R.P. *Writing - Original draft preparation*: Z.Z., C.R.P *Supervision*: C.R.P., M.S.L., R.T.B. *Writing-Reviewing and Editing*, Z.Z., T.S.H., M.S.L., R.T.B, C.R.P.

## Methods

### Animals and behavior

Six monkeys were implanted with microelectrode arrays (chronic). Two monkeys were implanted with 64 channels in three chronically implanted electrode arrays along the ventral stream (V1/V2, V4, posterior IT). One monkey was implanted with floating arrays (Microprobes for Life Sciences, Gaithsburg FL) in IT cortex. Three monkeys were implanted with Utah arrays (Blackrock Microsystems) in the right hemisphere operculum. The task for all monkeys was to maintain their gaze on a red fixation target (<0.2° radius) while large pictures were flashed at different locations on the screen. If the animals maintained their gaze within a 2.2° radius of the fixation target (1° for V1-array monkeys), they received a juice reward. All procedures received approval by Institutional Animal Care and Use Committees at Harvard Medical School and Washington University School of Medicine, procedures conformed to NIH guidelines provided in the Guide for the Care and Use of Laboratory Animals.

### Receptive field mapping

Each recording site’s receptive field was mapped by flashing a picture of a grayscale cartoon face in a 16°x16° grid at different locations relative to the fixation point with 2° spacing. The picture was flashed on for 100 ms and left off for 200 ms. The picture evoked firing rate responses from each array channel; responses were defined as the mean firing rate over 0-200 ms after picture onset minus the mean firing rate over the first 30 ms after picture onset. The subsequent 9 x 9 response matrices from every array channel were interpolated over 250 x 250 points. Each interpolated map was used to compute a receptive field, defined by regions where the mean firing rate exceeded the 99^th^ percentile of all responses in the map. The largest region exceeding this threshold was identified using the Matlab function *regionprops.m*, and its centroid location was used as the channel’s RF location. We defined the population RF as the mean array response. To characterize the reliability of individual RF estimates, we tested whether the site responded differentially to each position using a one-way ANOVA (picture position as level, *P* < 0.05 after false discovery rate correction). If the channel was reliable, we used the channel-specific RF estimate. Otherwise, we assigned it the population estimate. Since the RF center of multiunit sites in each array channel did not always overlap with the center of the heatmap, we averaged the RF location per channel, transformed it into a 2D vector in Cartesian space, and used it as a translation vector to shift the heatmap using the MATLAB function *imtranslate.m*.

### Stimuli

The stimuli were 16° x 16° natural-scene photographs embedded in a 30° x 30° brown-noise image. Each photo included natural (animals in the wild) and artificial settings (humans in laboratory settings) as well as computer-generated textures (bar fields). The pictures were presented in a 9° x 9° grid (spacing of 2°) with the center position located at the center of the population receptive field. The pictures were shown for 100-ms ON and 100 to 200-ms OFF, 3-14 flashes between juice rewards. Different pictures were interleaved within a trial. Every picture was presented at every position 30-37 times (V1 experiments) or 11-13 times (IT experiments). Each scene was later segmented independently using an automated semantic image segmentation model (DeepLab with ResNet-101 backbone^28^, trained on COCO-Stuff 164k dataset^29^) and a human-based segmentation task (Amazon Mechanical Turk, MTurk). All natural-scene photos were obtained from www.pexels.com and were free of copyright restrictions (one picture was provided by the Biomedical Primate Research Centre, Rijswijk, The Netherlands).

### Neuronal heatmaps

Heatmaps were evoked by randomly flashing each image of the natural scene at different positions relative to the center of the population RF, while the monkeys performed a fixation task. In their raw form, feature maps were matrices of average spike rates (200-ms-long, from 50 ms before image onset to 50 ms after image offset, 20 time-steps) computed per location (9 x 9), which could then be interpolated to the size of the input scene. The 9×9 maps were created by passing the spike rates and stimulus positions to *griddata.m*, then flipping the image left-right and up-down, as image coordinates have their lowest index value at the upper-left pixel, unlike regular numeric matrices that rely on Cartesian coordinates. For sites with RFs that were not centered at the fovea, an offset was added to the heatmap based on the eccentricity of the RF (measured independently of the heatmap experiments). All responses were based on a mix of single- and multiunits, though most channels were multiunits and visual hash. For population analyses, we computed each site’s heatmap normalized to the range 0–1, *H_j_a__*(*i*), and averaged across all:

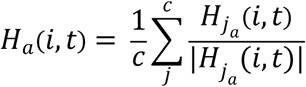

where *c* was the number of channels per array and *t* was the time step. To define a null hypothesis, for every time-averaged feature map, we scrambled the position information for every picture 999 times while keeping the array of responses the same across channels and across time.

### Semantic masks

Each scene used was segmented independently using both automated and human-based segmentation approaches. To get segmented scenes from DeepLab^28^, a ResNet101-based model, we adapted the unofficial PyTorch implementation and extracted labels from the COCO-Stuff 164k dataset^29^ within each scene. The pretrained weight *deeplabv2_resnet101_msc-cocostuff164k-100000* was downloaded from https://github.com/kazuto1011/deeplab-pytorch. Because there is not a *monkey* label in COCO-Stuff, and because DeepLab often classifies one object class (*tree*) with subordinate labels (*e.g., leaves*, *branch*), we merged the extracted labels into superordinate classes according to the COCO-Stuff label hierarchy. Ten *animal* labels and one *person* label were combined into a single *Animal* class; six *water* and fluid-related labels into a single *Water* class; nine *plant*-related labels such as *tree* and *moss* into a *Plant* class; eleven textile labels such as *blanket* and *rug* into a *Textile* class; four raw-material labels such as ‘*metal’* and ‘*plastic’* into a *Material* class; five structural and textual labels including *net* and *fence* into a *Structural* class; and six electronics labels such as *keyboard* into an E*lectronics* class. These classes were used as masks to segment the natural scenes.

In the human-guided approach, we first used Google Cloud Vision^11^ to provide ten labels that characterized each photograph, with the goal of asking human annotators to provide segmentation masks that best illustrated each label. We removed labels such as *organism* if they seemed too technical. After integration, there were 3–6 labels per scene. Participants on MTurk^30^ were asked to trace polygons around objects for each given label, each scene was randomly selected from the set of 36. Combinations of these maps were used to create higher-order categories such as “face,” “animacy,” and “environment.” There was a total of 27 possible semantic masks per scene. Each mask was defined by a binary matrix of the same size as the original natural scene.

To obtain the masks, we created a webpage for interactive semantic segmentation tasks on Amazon Mechanical Turk. The users were provided with the following information:

*Title: Trace polygons around the border of given labels (< 5 minutes)*

*Description: Apply a polygon over each object described per label in a precise and smooth way. The Task requires usually around 3 - 5 minutes*.

*Time Allotted: 8 Min*

*Qualifications Required: HIT approval rate (%) is greater than 7*

*Instructions:*

- This is a semantic segmentation task.
- To get started, first inspect the image. To the right, there is a list of labels indicating objects present in the image.

- Task is to apply a polygon over each object described per label.
- Click on any label in the box to activate that polygon. Left click to start annotating the image.
- Add points to make a polygon (mask) around the relevant region.
- When finished, press C to close the polygon.
- To undo, click Undo. To redo one label, click Delete to remove a selected polygon. Click Annotate to go back to annotating.
- It is okay to overlay the same image region with multiple labels if you find it appropriate.
- Please complete all labels in order to proceed to the next task.

All procedures received ethical approval by the Washington University Medical School Institutional Review Board.

To estimate how semantic masks contained neuronal or hidden unit activity, the following computations were performed:

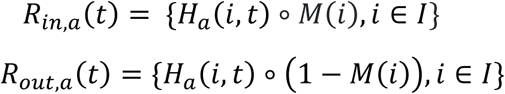

These plots led to a difference curve *D_a_*(*t*) = *R_in,a_*(*t*) − *R_out,a_* (*t*), and the area under the curve of *D_a_*(*t*) was used to summarize the overall activity. To control for the overall sparsity of the map, we shuffled the response regions for each scene to obtain a shuffled heatmap *H_a(shuffled)_*(*i, t*) and also measured the AUC for *D_a_shuffled__* (*t*).

### Computing feature-to-object ratio (FTO)

Objects were defined by semantic masks discussed in the previous section. The intuition behind the *feature-to-object ratio* (FTO) is to inspect the ratio of the localized feature over the semantic object in the heatmaps. There are two extreme cases: if the activation has the highest response only within the semantic object, then it results in a FTO of 1. On the contrary, if the responses all fell outside of the mask or were all zero, then FTO will be 0. Each unit-level heatmap was first normalized between 0 and 1. Then, we computed the normalized heatmap of array *a* that falls in the semantic mask *M_o_*(*i*) of object *o* and normalized by the semantic object size (FTO of 1 scenario). FTO was computed for each object in each array *a* in brain or layer in neural networks (with heatmap *H_a_*(*i*) replaced by a feature map *F_l_*(*i*))

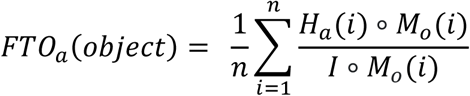

As a control, FTO was computed for each shuffled heatmap. We then fitted a linear regression line to the FTO(object) for Monkey A and Monkey B in V1/V2, V4, and IT and for each neural network in three selected layers discussed in detail below. The linear regression slope *k* for each *FTO*(*object*) across arrays or layers was estimated. For the statistical test, a one-way ANOVA test was performed between the *k*(*FTO*(*object*)) between objects and the p values were reported.

### Convolutional neural networks

We selected 14 different neural network models partially based on their ranking on Brain-Score^31^, going from 1–108. This set comprised *AlexNet*^32^, VGG-16^33^, CORNet-S and -Z^34^, ResNet-50^35^, ResNet-50-robust^36^, EfficientNet-B0^37^, VOneNet (ResNet-50 backbone, ResNet-50-robust backbone, ResNet 50 non-stochastic backbone, AlexNet backbone and CORNet-S backbone)^38^, DenseNet-201^39^, Xception^40^, and SqueezeNet^41^. In each network, we evenly selected three building blocks, identified the last convolutional layer per building block, and applied corresponding batch normalization and rectification to such convolutional layer. CORNet comprises four building blocks, each simulating the visual information processing of V1, V2, V4, or IT. We thus used the *V1nonlin*, *V2nonlin*, *V4nonlin*, and *ITnonlin* layers in CORNet-Z and *V1nonlin3*, *V2nonlin3*, *V4nonlin3*, and *ITnonlin3* layers in CORNet-S. AlexNet has five convolutional layers *conv1* to *conv5*, and we used all hidden units and applied rectification to these layers. VGG-16 has 16 weight layers and uses sequential 2-3 convolution-rectification layer motifs, which allows for a deeper trainable network. We used all units in the last convolutional layer of each motif: *conv1_2*, *conv3_3*, and *conv5_3.* In ResNet-50, there are five building blocks compressing repeating layer motifs, therefore we used layers *activation_1_relu*, *res2c_branch2c*, *res3d_branch2c*, *res4f_branch2c*, and *res5c_branch2c*, each coming from one building block, and with rectification applied accordingly. The VOneNet models comprise a VOne Block, simulating primate V1, followed by classic neural network backbones. Therefore, we used the *voneblock* with the convolutional layers in the last four blocks of the backbone network. For example, VOneNet ResNet-50 series all have a VOne Block followed by four ResNet building blocks, so we selected layers the same way as ResNet-50, except for replacing the first one by *voneblock*. Likewise, we chose the layers in VOneNet (ResNet-50 robust backbone), VOneNet (CORNet-S backbone), VOneNet (ResNet 50 non-stochastic backbone) following a similar procedure. DenseNet-201 is a 201-layer-deep, densely connected convolutional neural network and its convolution operations are distributed in one convolutional block followed by 4 dense blocks. We thus chose conv1*|relu*, *conv3_block12_2_conv*, *conv5_block32_2_conv*. EfficientNet B0 is a convolutional neural network baseline that is designed by neural architecture search. It has one convolution and seven mobile inverted bottleneck *MBConv* blocks followed by the final classification blocks. We selected the last convolutional layers from blocks *efficientnet-b0|blocks_0*, *efficientnet-b0|blocks_7*, and ‘*efficientnet-b0|blocks_15* and applied batch normalization and sigmoid rectification to each one of them. *Xception* uses deep wise separable convolutions in the network, and it has 36 convolutions spread out in 14 modules. Here we selected layer *block1_conv2_act*, *block7_sepconv3_act*, and *block14_sepconv3_act*. SqueezeNet comprises a standalone convolution block, eight Fire modules, and ended with a last convolutional block. We evenly chose three blocks in the architecture and used the last convolutional layer (after rectification) in each block: *fire2_relu_squeeze1×1, fire6_relu_squeeze1×1, and relu_conv10.* Every ANN hidden unit feature map was subsampled in an evenly spaced 9 x 9 grid and then normalized to the range of 0-1. Since ANNs do not have temporal responses in the activation, we simulated the temporal dynamics in ANNs (for comparison purpose) using normalized peristimulus time histogram (PSTH) from *in-vivo* experiments described above. Specifically, we multiplied the activation value from each hidden unit in the ANN with a randomly sampled PSTH. We randomly sampled V1 PSTH for hidden units in early layer, V4 PSTH for units in mid layer, and IT PSTH for those in the late layer.

### Comparison between models and brain

A number of statistical tests were conducted to compare the area under the curve (AUC) of *D* and the linear regression slope of *k* for each semantic object between regions (arrays *a* in brain or layers *l* in neural networks). Two sample Student’s t-test was performed between *D* in early and middle regions *D_early_*, *D_middle_* and in the middle and late regions *D_middle_* and *D_late_*. P values were reported. Spearman Correlation and Multivariate ANOVA (MANOVA) was conducted between *D_early_*, *D_middle_* and *D_late_* between monkeys and neural networks with reported p values. Spearman correlation tests whether the strength and direction of monotonic association between *D_early_*, *D_middle_* and *D_late_* between the monkey brain and neural networks are significant. MANOVA aims to test whether the mean vectors *D_block_* for each block in {*early, middle, late*} in brain or model are all equal to one another.

### Analyzing robustness in neural networks on parametric image degradations

We adapted the model vs human (https://github.com/bethgelab/model-vs-human) to evaluate the neural networks discussed above and tested on 17 datasets that can be downloaded from http://www.wichmannlab.org/. These datasets are associated with either parametric or binary distortions, such as color/grayscale alterations, contrast adjustments, or nonparametric image distortions such as texture-shape cue conflict. We evaluated the neural networks on all datasets and focused on the evaluation of texture-shape cue conflict. We added neural networks such as CORnet-S to the model zoo and data for each neural network was retrieved. For each data entry, the response was compared with both texture and shape categories to determine the correctness of classification. If the response did not match either the texture or shape, it was considered incorrect. Probabilities of correct texture-based classifications were calculated by dividing the count of correct texture-based classifications by the total count of corresponding combinations. Similarly, probabilities of correct shape-based classifications were calculated. Additionally, probabilities of incorrectly classifying texture instead of shape and vice versa were computed. We reported the probability of texture-based, shape-based, and incorrect classification for each neural network.

### Eye movement and saliency maps

In the gaze position experiments, the animals sat in front of the monitor. To start each trial, they acquired fixation on a center spot, after which a natural scene was presented. These scenes were the same stimuli as used in the neurophysiology experiments (30°x30° total size, with the central 18°x18° containing the photograph and the surround containing brown noise). The scenes were fixed to the center of the monitor. The animals were permitted to move their gaze anywhere within the scene within 1000 ms. To compute the eye fixation maps, the eye position in each trial was propagated into the ClusterFix toolbox^42^ to identify the first saccade after image onset. The eye position values (comprising a 1×2 tuple per ms) were binned into a 15×15 grid (15 bins were chosen for two reasons: the central canvas ratio was 18/30 = 0.6, so it was computationally convenient to have a grid size resulting in an integer when multiplied by 0.6, and also because the raw neuronal heatmaps measured 9×9 bins, and 15*0.6 = 9×9 fixation maps. These maps could then be interpolated to the original scene size, as the heatmaps were as well.

Saliency algorithms highlight the regions in visual scenes that are most likely to draw the attention of viewers, as measured perceptually or via saccades. We used three algorithms. The first was the Itti, Koch and Niebur algorithm^17^, which works by decomposing the visual image into feature channels using filter convolution (for orientation, color and intensity), implementing an antagonistic center-surround operation, normalizing each output, and then combining into a final saliency map. The second was the Graph-Based Visual Saliency model^11^, which works by extracting feature maps and using Markov chains over these maps, defining saliency as the equilibrium distribution over map regions. Both of these were applied using the code released under the *vision.caltech.edu* domain. The third was DeepGaze-II, which is based on Convolutional Neural Networks and adaptive gradient methods to estimate the gaze direction^18^.

