## supplementary tables for "Brain Feature Maps Reveal Progressive Animal-Feature Representations in the Ventral Stream"

### Supplementary Table A

*Monkey and ANN Median AUC of Response Difference and shuffled control for three areas*

| ***Model*** | **Layer** | **Median AUC of Response Difference** | **Shuffled AUC of Response Difference** |
|---|---|---|---|
| AlexNet | relu1 | 0.0659 ± 0.0705 | -0.1036 ± -0.0149 |
| | relu3 | -0.1209 ± -0.0155 | -0.0066 ± -0.0069 |
| | relu5 | 0.0355 ± 0.0332 | -0.0096 ± 0.0007 |
| *VGG-16* | relu1_1 | 0.0448 ± 0.0731 | 0.1139 ± 0.0496 |
| | relu3_3 | 0.2890 ± 0.1059 | -0.0805 ± -0.0063 |
| | relu5_3 | 0.1605 ± 0.0397 | -0.0070 ± 0.0079 |
| *ResNet-50* | activation_1_relu | 0.1975 ± 0.0769 | 0.0481 ± 0.0115 |
| | bn3d_branch2c | 0.1194 ± 0.0449 | 0.0064 ± 0.0007 |
| | bn5c_branch2c | 0.6012 ± 0.1472 | -0.0134 ± -0.0039 |
| *ResNet-50 robust* | activation_1_relu | 0.0076 ± 0.0395 | 0.0063 ± 0.0032 |
| | bn3d_branch2c | 0.1092 ± 0.0507 | 0.0251 ± 0.0063 |
| | bn5c_branch2c | 0.5409 ± 0.1020 | 0.0100 ± -0.0043 |
| *DenseNet-201* | conv1 | 0.0197 ± 0.0313 | -0.0056 ± -0.0063 |
| | conv3_block12_2_conv | 0.0558 ± 0.0315 | -0.0557 ± -0.0086 |
| | conv5_block32_2_conv | 0.3801 ± 0.1063 | -0.0175 ± 0.0023 |
| *EfficientNet-B0* | blocks_0 | -0.0698 ± -0.0288 | 0.0251 ± 0.0058 |
| | blocks_7 | -0.0699 ± -0.0272 | -0.0397 ± -0.0014 |
| | blocks_15 | 0.0462 ± -0.0090 | -0.0497 ± -0.0038 |
| *Xception* | block1_conv2_act | 0.1250 ± 0.0566 | 0.0350 ± 0.0044 |
| | block7_sepconv3_act | 0.0674 ± 0.0211 | 0.0335 ± 0.0032 |
| | block14_sepconv1_act | -0.0320 ± 0.0051 | -0.0576 ± 0.0001 |
| *SqueezeNet* | fire2-relu_squeeze1x1 | 0.1538 ± 0.0606 | 0.0212 ± 0.0050 |
| | fire6-relu_squeeze1x1 | 0.3727 ± 0.1089 | 0.0453 ± 0.0113 |
| | relu_conv10 | 0.8060 ± 0.1747 | 0.0672 ± 0.0264 |
| *CORNet-S* | V1nonlin2 | 0.0851 ± 0.0345 | 0.0319 ± 0.0027 |
| | V4nonlin3 | 0.2979 ± 0.0724 | -0.0400 ± -0.0034 |
| | ITnonlin3 | 0.7565 ± 0.1735 | -0.1291 ± -0.0127 |
| *CORNet-Z* | V1nonlin | 0.0839 ± 0.0358 | -0.0012 ± 0.0008 |
| | V4nonlin | 0.1076 ± 0.0493 | 0.0312 ± 0.0357 |
| | ITnonlin | 0.0437 ± 0.0553 | 0.0350 ± 0.0223 |
| *VOneNet_ResNet-50* | voneblock | 0.4743 ± 0.1124 | 0.0036 ± 0.0016 |
| | bn3d_branch2c | 0.0047 ± 0.0399 | 0.0456 ± 0.0022 |
| | bn5c_branch2c | 0.3951 ± 0.1030 | -0.0244 ± -0.0038 |
| *VOneNet_ResNet-50 (non stochastic)* | voneblock | 0.6031 ± 0.1380 | -0.0611 ± -0.0088 |
| | bn3d_branch2c | 0.0384 ± 0.0317 | -0.0101 ± 0.0021 |
| | bn5c_branch2c | 0.3041 ± 0.0620 | 0.0400 ± 0.0047 |
| *VOneNet_ResNet-50 robust* | voneblock | 0.4704 ± 0.1166 | -0.0534 ± -0.0023 |
| | bn3d_branch2c | 0.0575 ± 0.0284 | 0.0109 ± 0.0036 |
| | bn5c_branch2c | 0.6503 ± 0.1438 | 0.0541 ± 0.0138 |
| *VOneNet CORNet-S* | voneblock | 0.4555 ± 0.1170 | -0.0823 ± -0.0249 |
| | V4_nonlin3 | 0.1333 ± 0.0345 | -0.0207 ± -0.0025 |
| | IT_nonlin3 | 0.6111 ± 0.1543 | 0.0170 ± -0.0025 |
| *VOneNet AlexNet* | voneblock | 0.5557 ± 0.1145 | -0.0132 ± -0.0123 |
| | relu3 | -0.0762 ± 0.0030 | -0.0037 ± -0.0007 |

| | | | |
|---|---|---|---|
| | relu5 | 0.0348 ± 0.0206 | 0.0222 ± 0.0063 |
| *Monkeys (N=6)* | V1 | 0.0851 ± 0.0105 | -0.0092 ± -0.0004 |
| | V4 | 0.6322 ± 0.0248 | -0.0308 ± -0.0006 |
| | IT | 0.7414 ± 0.0940 | -0.0233 ± -0.0017 |
| *ViT-base* | block 0 | -0.5794 ± -0.1467 | 0.0274 ± 0.0308 |
| | block 6 | 2.1887 ± 0.4440 | 0.0276 ± 0.0122 |
| | block 11 | 1.1856 ± 0.2995 | -0.0958 ± -0.0093 |
| *ViT-small* | block 0 | -0.6512 ± -0.1149 | -0.0736 ± -0.0106 |
| | block 6 | 2.5125 ± 0.4690 | 0.0067 ± 0.0018 |
| | block 11 | 1.9548 ± 0.4247 | -0.1409 ± -0.0319 |
| *ViT-tiny* | block 0 | 0.3248 ± 0.0292 | -0.0719 ± -0.0031 |
| | block 6 | 0.1843 ± -0.0702 | 0.0302 ± -0.0030 |
| | block 11 | -0.2323 ± -0.0414 | 0.0419 ± 0.0094 |

### Supplementary Table B

*Linear model statistics of monkey and deep network's linear fit slope to variated objects*

| ***Model*** | **Animal features** | | | | | | **Non-animal features** | | | | | |
|---|---|---|---|---|---|---|---|---|---|---|---|---|
| | Primate face | Primate head | NP-head | NP-face | Body | Limbs | Food | Textile | Electronic s | Water | Other Art. Objects | Other Nat. objects |
| AlexNet | -0.4363 ± 0.1754 | -0.3144 ± 0.1464 | -0.1763 ± 0.2074 | -0.4948 ± 0.1709 | -0.0802 ± 0.1047 | -0.0872 ± 0.0476 | -0.2042 ± 0.1401 | 0.1151 ± 0.0782 | -0.0718 ± 0.0814 | 0.0544 ± 0.1967 | -0.2708 ± 0.0950 | -0.0887 ± 0.0781 |
| *VGG-16* | -0.4902 ± 0.1985 | -0.3873 ± 0.1755 | -0.1125 ± 0.2262 | -0.3459 ± 0.1507 | -0.0576 ± 0.1285 | 0.0054 ± 0.0576 | -0.0371 ± 0.2059 | 0.1209 ± 0.1149 | -0.0615 ± 0.0746 | 0.1379 ± 0.1806 | -0.2409 ± 0.1053 | -0.0824 ± 0.0995 |
| *ResNet-50* | 0.2693 ± 0.1807 | 0.1614 ± 0.1433 | 0.1386 ± 0.2081 | 0.1370 ± 0.1745 | 0.2622 ± 0.0986 | 0.2385 ± 0.0584 | 0.2674 ± 0.1513 | -0.1530 ± 0.0794 | 0.0893 ± 0.0945 | -0.0392 ± 0.0858 | -0.1413 ± 0.0707 | -0.2534 ± 0.0662 |
| *DenseNet-201* | 0.2431 ± 0.1292 | 0.1929 ± 0.1177 | 0.1070 ± 0.1442 | 0.1495 ± 0.1046 | 0.1741 ± 0.0782 | 0.1505 ± 0.0490 | 0.2062 ± 0.1433 | 0.0592 ± 0.0934 | 0.2075 ± 0.0712 | -0.0335 ± 0.1320 | 0.0190 ± 0.0885 | -0.0016 ± 0.0877 |
| *EfficientNet-B0* | 0.0717 ± 0.1293 | 0.2091 ± 0.1107 | 0.3136 ± 0.1468 | 0.3310 ± 0.1273 | 0.1017 ± 0.0661 | 0.1149 ± 0.0444 | 0.1047 ± 0.1362 | -0.0144 ± 0.0642 | 0.0500 ± 0.0626 | -0.1711 ± 0.1416 | -0.0360 ± 0.0891 | -0.0839 ± 0.0728 |
| *Xception* | -0.2503 ± 0.1299 | -0.1954 ± 0.1333 | -0.3948 ± 0.1583 | -0.4024 ± 0.1138 | -0.1500 ± 0.0756 | -0.0809 ± 0.0388 | -0.1965 ± 0.0983 | 0.0059 ± 0.0623 | -0.0672 ± 0.0629 | 0.0473 ± 0.0613 | -0.1759 ± 0.0632 | -0.1009 ± 0.0520 |
| *SqueezeNet* | 0.4107 ± 0.1524 | 0.3565 ± 0.1449 | 0.4558 ± 0.1688 | 0.4617 ± 0.1493 | 0.1966 ± 0.0976 | 0.1814 ± 0.0696 | 0.4355 ± 0.2328 | 0.1875 ± 0.1244 | 0.0116 ± 0.1226 | -0.0266 ± 0.1146 | -0.0613 ± 0.0987 | -0.3801 ± 0.1031 |
| *CORNet-S* | 0.5403 +- 0.1223 | 0.3377 +- 0.0960 | 0.4071 +- 0.1374 | 0.2328 +- 0.0951 | 0.3158 +- 0.0696 | 0.2285 +- 0.0401 | 0.5069 +- 0.1581 | 0.0392 +- 0.0952 | 0.0933 +- 0.0935 | -0.2135 +- 0.1484 | 0.0100 +- 0.0759 | -0.1113 +- 0.0653 |
| *CORNet-Z* | 0.0740 ± 0.1913 | 0.1328 ± 0.1633 | 0.1798 ± 0.1454 | -0.1098 ± 0.2051 | 0.0469 ± 0.1013 | 0.0798 ± 0.0650 | -0.4543 ± 0.2097 | -0.1500 ± 0.1519 | -0.3606 ± 0.1075 | -0.1131 ± 0.1443 | -0.1200 ± 0.0776 | -0.1121 ± 0.0490 |
| *VOneNet_ResNet-50* | -0.1365 ± 0.1831 | -0.1793 ± 0.1562 | -0.0616 ± 0.2590 | -0.2893 ± 0.1667 | -0.0113 ± 0.1111 | -0.0460 ± 0.0710 | -0.2254 ± 0.1837 | 0.1466 ± 0.1071 | -0.0668 ± 0.0848 | 0.2426 ± 0.1512 | -0.1987 ± 0.0826 | -0.1782 ± 0.0788 |
| *VOneNet_ResNet-50 (non stochastic)* | -0.3653 ± 0.2025 | -0.4460 ± 0.1489 | -0.1850 ± 0.2122 | -0.3476 ± 0.2259 | -0.2001 ± 0.1136 | -0.2183 ± 0.0722 | -0.4963 ± 0.1818 | 0.0671 ± 0.1237 | 0.0011 ± 0.0979 | 0.3054 ± 0.1575 | -0.1358 ± 0.1066 | -0.1487 ± 0.0998 |
| *VOneNet_ResNet-50 robust* | 0.2206 ± 0.1928 | 0.0364 ± 0.1712 | 0.1067 ± 0.2301 | -0.0201 ± 0.2601 | 0.0699 ± 0.1121 | 0.1252 ± 0.0734 | 0.1748 ± 0.2242 | 0.0403 ± 0.1294 | 0.0490 ± 0.1148 | 0.1791 ± 0.1308 | -0.0896 ± 0.1117 | -0.2358 ± 0.0876 |
| *VOneNet CORNet-S* | 0.0745 ± 0.1692 | -0.1203 ± 0.1549 | 0.1532 ± 0.2060 | -0.2866 ± 0.1948 | 0.1466 ± 0.1153 | 0.0855 ± 0.0711 | -0.2445 ± 0.1721 | 0.0792 ± 0.1262 | -0.0161 ± 0.0898 | 0.2177 ± 0.1832 | -0.1296 ± 0.0920 | -0.2366 ± 0.0892 |
| *VOneNet AlexNet* | -0.4172 ± 0.1426 | -0.4162 ± 0.1276 | -0.4230 ± 0.2155 | -0.4571 ± 0.1809 | -0.2659 ± 0.0827 | -0.2047 ± 0.0549 | -0.3863 ± 0.1434 | 0.2394 ± 0.0976 | -0.1043 ± 0.0685 | 0.2139 ± 0.1281 | -0.2574 ± 0.0871 | -0.1017 ± 0.0667 |
| *Monkeys (N=6)* | 0.7140 ± 0.0750 | 0.5920 ± 0.0670 | 0.5306 ± 0.1052 | 0.4289 ± 0.0992 | 0.3872 ± 0.0440 | 0.2521 ± 0.0283 | 0.1695 ± 0.0679 | 0.1616 ± 0.0507 | 0.1231 ± 0.0506 | 0.0260 ± 0.0884 | -0.0043 ± 0.0654 | -0.0982 ± 0.0716 |
| *ViT-tiny* | 0.0812 ± 0.2340 | 0.0930 ± 0.1964 | 0.0247 ± 0.3383 | -0.0371 ± 0.1967 | -0.2442 ± 0.1744 | -0.0550 ± 0.1014 | -0.6809 ± 0.2659 | -0.3751 ± 0.1081 | -0.1827 ± 0.1868 | 0.6102 ± 0.3939 | 0.7423 ± 0.1481 | 0.2157 ± 0.2005 |
| *ViT-small* | 2.2459 ± 0.4356 | 1.9002 ± 0.3741 | 2.3331 ± 0.6354 | 2.5923 ± 0.5383 | 1.2703 ± 0.2462 | 1.0847 ± 0.1685 | 1.4079 ± 0.4413 | -0.6598 ± 0.2771 | 0.4055 ± 0.2011 | -0.6672 ± 0.2431 | 0.6562 ± 0.2415 | -0.2991 ± 0.2059 |
| *ViT-base* | 2.2132 ± 0.4011 | 1.7643 ± 0.3627 | 1.6815 ± 0.6127 | 2.1373 ± 0.5243 | 1.0308 ± 0.2372 | 0.8420 ± 0.1555 | 1.7466 ± 0.4012 | -0.4575 ± 0.2261 | 0.4581 ± 0.1961 | -0.3779 ± 0.3215 | 0.6413 ± 0.2125 | -0.1388 ± 0.2184 |

### Supplementary Table C

*Statistical tests of monkey and deep network's linear fit slope to animal and non-animal feature*

| Model | Animal features | | | | | Non-animal features | | | |
|---|---|---|---|---|---|---|---|---|---|
| | $k_a$ (median ± SE) | two sample t-test p value ($D_{l_{early}}$, $D_{l_{middle}}$) | two sample t-test p value ($D_{l_{middle}}$, $D_{l_{late}}$) | MANOVA p value | Spearman Correlation $\rho$ (p value) | $k_a$ (median ± SE) | two sample t-test p value ($D_{l_{early}}$, $D_{l_{middle}}$) | two sample t-test p value ($D_{l_{middle}}$, $D_{l_{late}}$) | Spearman Correlation $\rho$ (p value) |
| AlexNet | -0.216±0.198 | 0.041 | 0.006 | 0.005 | 0.013 (9.0e-01) | -0.262±0.069 | 0.001 | 0.865 | -0.179 (1.3e-03) |
| VGG-16 | -0.198±0.240 | 0.672 | 0.060 | 0.027 | 0.078 (4.7e-01) | -0.224±0.084 | 0.122 | 0.316 | -0.149 (7.9e-03) |
| ResNet-50 | 0.351±0.194 | 0.270 | 0.001 | 0.006 | 0.370 (3.3e-04) | -0.198±0.067 | <0.0001 | 0.894 | -0.196 (4.5e-04) |
| ResNet-50-robust | 0.334±0.143 | 0.843 | 0.006 | 1.19E-36 | 0.435 (1.8e-05) | 0.097±0.061 | 0.828 | 0.074 | 0.078 (1.7e-01) |
| DenseNet-201 | 0.388±0.135 | 0.812 | 0.003 | 0.401 | 0.362 (4.5e-04) | 0.093±0.068 | 0.001 | <0.0001 | 0.120 (3.2e-02) |
| EfficientNet-B0 | 0.141±0.131 | 0.744 | 0.432 | 0.095 | 0.092 (3.9e-01) | 0.048±0.058 | 0.858 | 0.416 | 0.043 (4.4e-01) |
| Xception | -0.281±0.164 | 0.277 | 0.474 | 0.045 | -0.174 (1.0e-01) | -0.212±0.045 | 0.001 | 0.206 | -0.205 (2.3e-04) |
| SqueezeNet | 0.589±0.176 | 0.223 | 0.036 | 1.11E-05 | 0.465 (4.0e-06) | -0.187±0.099 | 0.028 | 0.775 | -0.186 (8.8e-04) |
| CORnet-S | 0.750±0.141 | 0.136 | <0.0001 | 4.87E-15 | 0.532 (7.0e-08) | 0.091±0.068 | 0.222 | 0.045 | -0.005 (9.2e-01) |
| CORnet-Z | 0.104±0.201 | 0.809 | 0.712 | 0.805 | 0.066 (5.4e-01) | -0.363±0.072 | <0.0001 | 0.663 | -0.246 (8.7e-06) |
| VOneNet_ResNet-50 | -0.106±0.211 | 0.045 | 0.034 | 3.16E-21 | 0.003 (9.8e-01) | -0.254±0.078 | 0.009 | 0.501 | -0.158 (4.7e-03) |
| VOneNet_ResNet-50(non stochastic) | -0.443±0.217 | 0.009 | 0.263 | 0.002 | -0.218 (3.9e-02) | -0.202±0.088 | 0.001 | 0.115 | -0.069 (2.2e-01) |
| VOneNet_ResNet-50 robust | 0.095±0.204 | 0.006 | <0.0001 | 0.773 | 0.106 (3.2e-01) | -0.120±0.092 | 0.018 | 0.361 | -0.080 (1.5e-01) |
| VOneNet-CORnetS | 0.164±0.191 | 0.010 | <0.0001 | 0.469 | 0.186 (7.8e-02) | -0.169±0.080 | 0.042 | 0.938 | -0.087 (1.2e-01) |
| VOneNet-AlexNet | -0.538±0.159 | 0.001 | 0.200 | 0.775 | -0.303 (3.7e-03) | -0.257±0.065 | 0.004 | 0.232 | -0.176 (1.6e-03) |
| ViT-base | 2.428±0.422 | <0.0001 | 0.075 | 0.045 | 0.523 (1.2e-07) | 0.717±0.180 | 0.001 | 0.541 | 0.227 (4.2e-05) |

| | | | | | | | | | |
|---|---|---|---|---|---|---|---|---|---|
| ViT-small | 2.952±0.400 | <0.0001 | 0.847 | 1.49E-30 | 0.672 (4.1e-13) | 0.513±0.195 | 0.026 | 0.662 | 0.150 (7.4e-03) |
| ViT-tiny | -0.392±0.309 | 0.179 | 0.765 | 0.018 | -0.196 (6.4e-02) | 0.256±0.134 | 0.336 | 0.297 | 0.062 (2.7e-01) |
| Monkeys (N=6) | 0.384±0.040 | <0.0001 | 0.057 | N/A | 0.539 (9.2e-23) | 0.005±0.024 | 0.014 | 0.017 | 0.008 (8.1e-01) |

**Supplementary Table D**

*FTO median±SE and median k ± SE for fitting FTO in various object classes (compared to control value ± SE)*

| | *Head* | | *Animal* | | *Natural Texture* | |
|---|---|---|---|---|---|---|
| Monkey A | 0.55 ± 0.00029<br>0.59 ± 0.00036<br>0.62 ± 0.00019<br><br>(0.55 ± 0.00024<br>0.49 ± 0.00033<br>0.49 ± 0.00019) | 0.034±0.004<br>(-0.028±0.004) | 0.51 ± 0.00019<br>0.53 ± 0.00023<br>0.55 ± 0.00013<br><br>(0.52 ± 0.00017<br>0.48 ± 0.00020<br>0.46 ± 0.00012) | 0.011±0.003<br>(-0.037±0.003) | 0.55 ± 0.00074<br>0.47 ± 0.0010<br>0.40 ± 0.00051<br><br>(0.55 ± 0.00069<br>0.46 ± 0.00095<br>0.41 ± 0.00049) | -0.071±0.006<br>(-0.073±0.006) |
| Monkey B | 0.47 ± 0.00037<br>0.53 ± 0.00046<br>0.62 ± 0.00041<br><br>(0.50 ± 0.00031<br>0.44 ± 0.00039<br>0.48 ± 0.00034) | 0.057±0.006<br>(-0.012±0.005) | 0.45 ± 0.00024<br>0.48 ± 0.00031<br>0.53 ± 0.00027<br><br>(0.49 ± 0.00020<br>0.44 ± 0.00026<br>0.45 ± 0.00023) | 0.032±0.005<br>(-0.026±0.004) | 0.47 ± 0.00087<br>0.40 ± 0.0011<br>0.48 ± 0.00098<br><br>(0.49 ± 0.00073<br>0.40 ± 0.0011<br>0.45 ± 0.00095) | -0.011±0.007<br>(-0.027±0.007) |
| ANOVA | F=30.014 p=1.5e-13(df=2,1715)<br>F=27.640 p=1.5e-12(df=2,1715)<br><br>F=43.047 p=7.6e-19(df=2,1342)<br>F=15.717 p=1.8e-07(df=2,1342) | F=21.00266 $p=4.5\times10^{-2}$(df=1,2) | F=5.713 p=3.4e-03(df=2,2169)<br>F=80.596 p=1.7e-34(df=2,2169)<br><br>F=25.305 p=1.5e-11(df=2,1747)<br>F=40.859 p=4.5e-18(df=2,1747) | F= 20.36575 $p=4.6\times10^{-2}$(df=1,2) | F=69.458 p=5.7e-27(df=2,444)<br>F=82.497 p=3.4e-31(df=2,444)<br><br>F=4.510 p=1.2e-02(df=2,337)<br>F=14.605 p=8.3e-07(df=2,337) | F=0.06427 $p=8.2\times10^{-1}$ (df=1,2) |

### Supplementary Table E

*Performance (accuracy) of selected ANNs to adversarial attacks*

| Dataset name | CORNet-S | AlexNet | VGG-16 | ResNet-50 | ResNet-50_l2_eps0 | ResNet-50_l2_eps0_5 | ResNet-50_l2_eps1 | ResNet-50_l2_eps3 | ResNet-50_l2_eps5 |
|---|---|---|---|---|---|---|---|---|---|
| Stylized (top 1) | 0.25 | 0.27875 | 0.295 | 0.1375 | 0.34375 | 0.4375 | 0.44125 | 0.4175 | 0.39 |
| Stylized (top 5) | 0.4625 | 0.6175 | 0.62625 | 0.54375 | 0.6775 | 0.75375 | 0.74 | 0.73 | 0.6875 |
| edge (top 1) | 0.177344 | 0.29375 | 0.21875 | 0.18203125 | 0.2375 | 0.225 | 0.20625 | 0.25625 | 0.275 |
| Silhouette (top 1) | 0.973438 | 0.43125 | 0.475 | 0.97578125 | 0.48125 | 0.55 | 0.6 | 0.60625 | 0.6 |
| cue-conflict (top 1) | 0.636719 | 0.1890625 | 0.145313 | 0.68046875 | 0.174219 | 0.295313 | 0.332031 | 0.439844 | 0.474219 |
| Colour (top 1) | 0.335156 | 0.909375 | 0.967969 | 0.325 | 0.978125 | 0.980469 | 0.969531 | 0.945313 | 0.897656 |
| Contrast (top 1) | 0.397656 | 0.396875 | 0.648438 | 0.4125 | 0.701563 | 0.454688 | 0.401563 | 0.335156 | 0.299219 |
| high-pass (top 1) | 0.502679 | 0.29375 | 0.351563 | 0.508928571 | 0.3375 | 0.289063 | 0.286719 | 0.261719 | 0.263281 |
| low-pass (top 1) | 0.8375 | 0.33515625 | 0.372656 | 0.845535714 | 0.389844 | 0.414063 | 0.415625 | 0.403906 | 0.365625 |
| phase-scrambling (top 1) | 0.94375 | 0.444642857 | 0.484821 | 0.960714286 | 0.491964 | 0.519643 | 0.522321 | 0.538393 | 0.496429 |
| power-equalisation (top 1) | 0.732031 | 0.684821429 | 0.816964 | 0.7546875 | 0.836607 | 0.842857 | 0.826786 | 0.778571 | 0.725893 |
| false-colour (top 1) | 0.471875 | 0.866964286 | 0.951786 | 0.47421875 | 0.95625 | 0.961607 | 0.934821 | 0.907143 | 0.877679 |
| Rotation (top 1) | 0.407031 | 0.57109375 | 0.742969 | 0.4234375 | 0.739063 | 0.723438 | 0.708594 | 0.6375 | 0.547656 |
| Eidolon I (top 1) | 0.346094 | 0.415625 | 0.44375 | 0.37109375 | 0.444531 | 0.51875 | 0.536719 | 0.546094 | 0.535156 |
| Eidolon II (top 1) | 0.364844 | 0.34921875 | 0.38125 | 0.40546875 | 0.403125 | 0.421094 | 0.420313 | 0.417188 | 0.403125 |
| Eidolon III (top 1) | 0.6 | 0.3 | 0.325 | 0.59625 | 0.338281 | 0.369531 | 0.386719 | 0.388281 | 0.371094 |
| uniform-noise (top 1) | 0.79375 | 0.20625 | 0.399219 | 0.8025 | 0.379688 | 0.341406 | 0.316406 | 0.260156 | 0.234375 |

| | | | | | | | | | |
|---|---|---|---|---|---|---|---|---|---|
| Sketch (top 1) | 0.3625 | 0.45 | 0.57625 | 0.37125 | 0.61375 | 0.6425 | 0.63375 | 0.59375 | 0.555 |
| Sketch (top 5) | 0.67875 | 0.71625 | 0.785 | 0.695 | 0.81 | 0.8075 | 0.82875 | 0.795 | 0.75125 |

**Supplementary Table E *(continued)***

*Performance (accuracy) of selected ANNs to adversarial attacks*

| Dataset name | Densenet-201 | SqueezeNet | EfficientNet-B0 | CORNe-Z | Vonenet_AlexNet | Vonenet CORNet-S | Vonenet ResNet-50 | Vonenet ResNet-50 (non stochastic) | Vonenet ResNet-50 robust |
|---|---|---|---|---|---|---|---|---|---|
| Stylized (top 1) | 0.41875 | 0.25375 | 0.475 | 0.24 | 0.2425 | 0.15125 | 0.13125 | 0.10375 | 0.3575 |
| Stylized (top 5) | 0.7425 | 0.5975 | 0.77875 | 0.58625 | 0.56625 | 0.445 | 0.4425 | 0.37375 | 0.66875 |
| edge (top 1) | 0.38125 | 0.15 | 0.3625 | 0.15625 | 0.20625 | 0.0625 | 0.0625 | 0.0625 | 0.23125 |
| Silhouette (top 1) | 0.5125 | 0.24375 | 0.53125 | 0.2125 | 0.25 | 0.0625 | 0.0625 | 0.0625 | 0.425 |
| cue-conflict (top 1) | 0.211719 | 0.135156 | 0.209375 | 0.136719 | 0.242969 | 0.141406 | 0.119531 | 0.095313 | 0.361719 |
| Colour (top 1) | 0.982813 | 0.878125 | 0.982813 | 0.825781 | 0.809375 | 0.463281 | 0.380469 | 0.275 | 0.929688 |
| Contrast (top 1) | 0.807031 | 0.460938 | 0.733594 | 0.330469 | 0.382031 | 0.520313 | 0.532031 | 0.655469 | 0.48125 |
| high-pass (top 1) | 0.465625 | 0.33125 | 0.342188 | 0.305469 | 0.260156 | 0.240625 | 0.211719 | 0.214063 | 0.305469 |
| low-pass (top 1) | 0.442969 | 0.308594 | 0.414844 | 0.267969 | 0.274219 | 0.176563 | 0.160156 | 0.1375 | 0.391406 |
| phase-scrambling (top 1) | 0.546429 | 0.40625 | 0.496429 | 0.372321 | 0.383036 | 0.191071 | 0.170536 | 0.150893 | 0.501786 |
| power-equalisation (top 1) | 0.904464 | 0.654464 | 0.857143 | 0.553571 | 0.554464 | 0.283929 | 0.248214 | 0.182143 | 0.791964 |
| false-colour (top 1) | 0.975893 | 0.857143 | 0.974107 | 0.772321 | 0.775 | 0.397321 | 0.353571 | 0.225 | 0.905357 |
| Rotation (top 1) | 0.790625 | 0.595313 | 0.794531 | 0.519531 | 0.427344 | 0.23125 | 0.2 | 0.174219 | 0.578125 |
| Eidolon I (top 1) | 0.465625 | 0.349219 | 0.457813 | 0.338281 | 0.39375 | 0.232031 | 0.182031 | 0.142188 | 0.492969 |
| Eidolon II (top 1) | 0.419531 | 0.308594 | 0.414844 | 0.296875 | 0.3 | 0.182813 | 0.142969 | 0.120313 | 0.395313 |
| Eidolon III (top 1) | 0.360156 | 0.270313 | 0.373438 | 0.255469 | 0.26875 | 0.157813 | 0.135156 | 0.114844 | 0.371094 |
| uniform-noise (top 1) | 0.483594 | 0.232813 | 0.442969 | 0.18125 | 0.275781 | 0.41875 | 0.425781 | 0.435156 | 0.409375 |
| Sketch (top 1) | 0.6775 | 0.44375 | 0.63625 | 0.4175 | 0.3175 | 0.075 | 0.075 | 0.065 | 0.49 |
| Sketch (top 5) | 0.855 | 0.71 | 0.84375 | 0.72125 | 0.585 | 0.3625 | 0.33125 | 0.32875 | 0.7175 |
